# Is a Ring-arranged Vascular Pattern Unique to Stems? A Three Dimensional Reconstruction of *Anaxagorea* (Annonaceae) Carpel Vasculature and its Evolutionary Implications

**DOI:** 10.1101/2020.05.22.111716

**Authors:** Ya Li, Wei Du, Ye Chen, Shuai Wang, Xiao-Fan Wang

**Affiliations:** College of Life Sciences, Wuhan University, Wuhan, Hubei 430072, China; Department of Environmental Art Design, Tianjin Arts and Crafts Vocational College, Tianjin, Tianjin 300250, China; College of Life Sciences and Environment, Hengyang Normal University, Hengyang, Hunan 421001, China

**Keywords:** 3D reconstruction, *Anaxagorea*, angiosperms, organogenesis, origin of the carpel, vascular anatomy

## Abstract

Elucidating the origin of carpel has been a challenge in botany for a long time. The Unifying Theory suggested that the carpel originate from a composite organ comprising an ovule-bearing shoot and a foliar part enclosing the shoot. A logical inference from this theory is that placenta in angiosperms should have radiosymmetrical vasculature, just like that in a young branch. *Anaxagorea* is the most basal genus of the primitive angiosperm family, Annonaceae. The conspicuous carpel stipe makes it an ideal material for exploring the carpel vasculature. In this study, serial sections of flower and carpel were delineated in *Anaxagorea luzonensis* and *A. javanica*, and a three-dimensional model of the carpel vasculature was reconstructed. The results show that (1) vascular bundles at both the carpel stipe and the ovule/placenta are in a radiosymmetrical pattern, (2) the amphicribral bundles would develop into ring-arranged bundle complex with the carpel maturation, (3) the ovule/placenta bundles were separated from the bundles of the carpel wall, and, (4) all the radiosymmetrical vasculature (including amphicribral bundles and ring-arranged bundle complexes) in the carpel were fed by a larger radiosymmetrical bundle system. These results suggest that the radiosymmetrical pattern of carpel vasculature are in line with the Unifying Theory.

## INTRODUCTION

Based on common textbook knowledge (Esau, 1965; Rudall, 2007; Evert & Eichhorn, 2013), the arrangement of vascular bundles —in a radiosymmetrical pattern or lateral pattern — is an important difference between the axial organ (such as stems), and the lateral organ (such as leaves). Amphicribral bundles are frequently encountered in stems of early land plants (Ogura, 1972; Fahn, 1990). An amphicribral bundle is featured by xylem surrounded by phloem, while a collateral bundle has adaxial xylem, and abaxial phloem. De Bary (1884) observed that amphicribral vascular bundles are apparently rare in angiosperms. However, their occurrence has been reported in flowers, especially in the placenta and the ovules of the carpel. They are fed independently from lateral bundles which supply the carpel wall (e.g., *Actinidia* [Guo et al., 2013], *Magnolia* [Liu et al., 2014]; *Michelia* [Zhang et al., 2017]; *Dianthus* [Guo et al., 2017]; *Tapiscia* [Xin et al., 2019]). These studies, based on comparison of vascular bundles, suggest that the widespread existence of amphicribral bundles in the ovule/placenta is compatible with the Unifying Theory (Wang, 2010, 2018), which states that the carpel in the classic sense is a composite organ comprising an ovule-bearing shoot (ovule/placenta) and a foliar part (carpel wall) enclosing the shoot.

Although vascular bundles are generally considered conservative, their anatomy is influenced by the immediately adjacent histological environment, and its histology is greatly influenced by the next older vascular bundle to which it connects (Endress, 2019). Could the carpel vasculature provide threads for speculate the evolutionary origin of carpels? In the past, the material usually included limited developmental stages, either flower or fruit. To account for the different development stages of vascular bundles and the adjacent histological environment, two species of *Anaxagorea* were selected from both flowering and fruiting stages, and continuous serial sections were conducted of their receptacles and carpel, and a three dimensionl (3D) -model of the carpel vasculature was reconstructed.

*Anaxagorea* is the most basal genus of Annonaceae, one of the early radiations of angiosperms lineage (Doyle & Le Thomas, 1996; Doyle et al., 2004; Chatrou et al., 2012; Chatrou et al., 2018). *Anaxagorea* carpels are apocarpous throughout their life history (Deroin, 1988), and each has a notably long carpel stipe (Endress &Armstrong, 2011). If the Unifying Theory of carpel origin is practical, then it should also be applicable for *Anaxagorea*, i.e., (1) the ovule/placenta bundle of *Anaxagorea* should be amphicribral, (2) the ovule/placenta bundle should always be separated from the vascular bundle of the carpel wall, and hence, (3) the amphicribral bundle should come from a larger axial bundle system.

## MATERIALS AND METHODS

### Scanning electron microscopy and paraffin sectioning

*Anaxagorea luzonensis* flower samples at different floral stages (from early bud to young fruit) were collected from the Diaoluo Mountain, Hainan, China, in July 2017 and *Anaxagorea javanica* from the Xishuangbanna Tropical Botanical Garden, Yunnan, China in May 2017. The gynoecia were isolated and preserved in 70% formalin-acetic acid-alcohol (5:5:90, v/v), and the fixed specimens were dehydrated in a 50% to 100% alcohol series. To delineate the structure and development of the carpel, carpels were removed from the gynoecia, passed through an iso-pentanol acetate series (SCR, Shanghai, China), critically point-dried, sputter-coated with gold, observed, and photographed under a scanning electron microscope (Tescan VEGA-3-LMU, Brno, Czech Republic). Flowers and carpels were embedded in paraffin, serially sectioned into 10–12-µm thick sections, and stained with Safranin O and Fast Green to illustrate the vasculature. The transverse sections were examined and photographed using a bright-field microscope (Olympus BX-43-U, Tokyo, Japan). In addition, longitudinal hand-cut sections were made and observed for a rough check and better understanding of the vasculature.

### Topological analysis of carpel vasculature

Consecutive paraffin sections, 12-µm each, of *A. javanica* were stained with aniline blue, examined and photographed after excitation at 365 nm using an epifluorescence microscope (Olympus BX-43-U, Tokyo, Japan) and a semiconductor refrigeration charged coupled device (RisingCam, MTR3CMOS). Manual image registration of each dataset was arried outusing Photoshop CC 2017 (Adobe, San Jose, CA, US). Forty-five images were selected equidistant from the 423 sections taken for the 3D reconstruction. The figures were organized according to the vascular bundle outlines of the sections by using Photoshop CC 2017 and Illustrator CC 2017 (Adobe). The xylem and phloem contours were manually drawn, extracted as paths with the pen tool, and exported in DWG format. The DWG files were imported into 3Ds Max 2016 (Autodesk, San Rafael, CA, US) and sorted according to the distance and order of the sections. The paths were converted to Editable Spline curves to generate the basic modeling contour. The Loft command of Compound Objects was used to get the shape of the Editable Spline, and a complete 3D carpel vasculature model was generated.

## RESULTS

### Gynoecium structure and carpel organogenesis

Flowers of two study species were trimerous with a whorl of sepals, two morphologically distinct whorls of petals, and numerous stamens (and inner staminodes of *A. Javanica*) (Figs. 1A–1D). *A. luzonensis* usually exhibits two to four completely separate carpels (Figs. 1A, 1G). The carpel primordia are almost hemispherically initiated and larger than the stamen primordia (Fig. 1F). Each carpel consists of a plicate zone, a very short ascidiate zone (Figs. 3G, 5I, 5J), and a long, conspicuous stipe (Fig. 2F). Carpel stipe ontogenesis occurs at the early stages of carpel development (Fig. 2B). The continuous growth of the flanks on the ventral side of the young carpel triggers its early closure; however, the closure does not extend to the base of the carpel, where the carpel stipe was previously present (Fig. 2C). Subsequently, the dorsal region of each carpel thickens markedly, and the stigma forms (Figs. 2D, 2E). At anthesis, the carpels are widest at the basal region with an arch on the abaxial side. The carpel stipe remains elongate, accounting for approximately a quarter of the carpel length at anthesis, and continues to elongate during the fruiting stage (Fig. 1F). Each carpel has two lateral ovules with the placentae at the ovary base (Figs. 3H, 5L). *A. Javanica* exhibits a multicarpellate gynoecium (Figs. 1B, 1J). The carpels are completely separate and appear whorled at initiation (Fig. 1I); as the carpel volume increases, the whorled structure becomes less obvious because the space in floral apex becomes limited. Each carpel consists of a plicate zone and a conspicuous carpel stipe (Fig. 2J) but lacks the short ascidiate zone. The carpel stipe ontogenesis occurs in the early stages of carpel development (Fig. 2H) and remains elongate during the flowering and fruiting stages (Figs. 1D, 2I–2J). Each carpel has two lateral ovules.

**Fig. 1.**
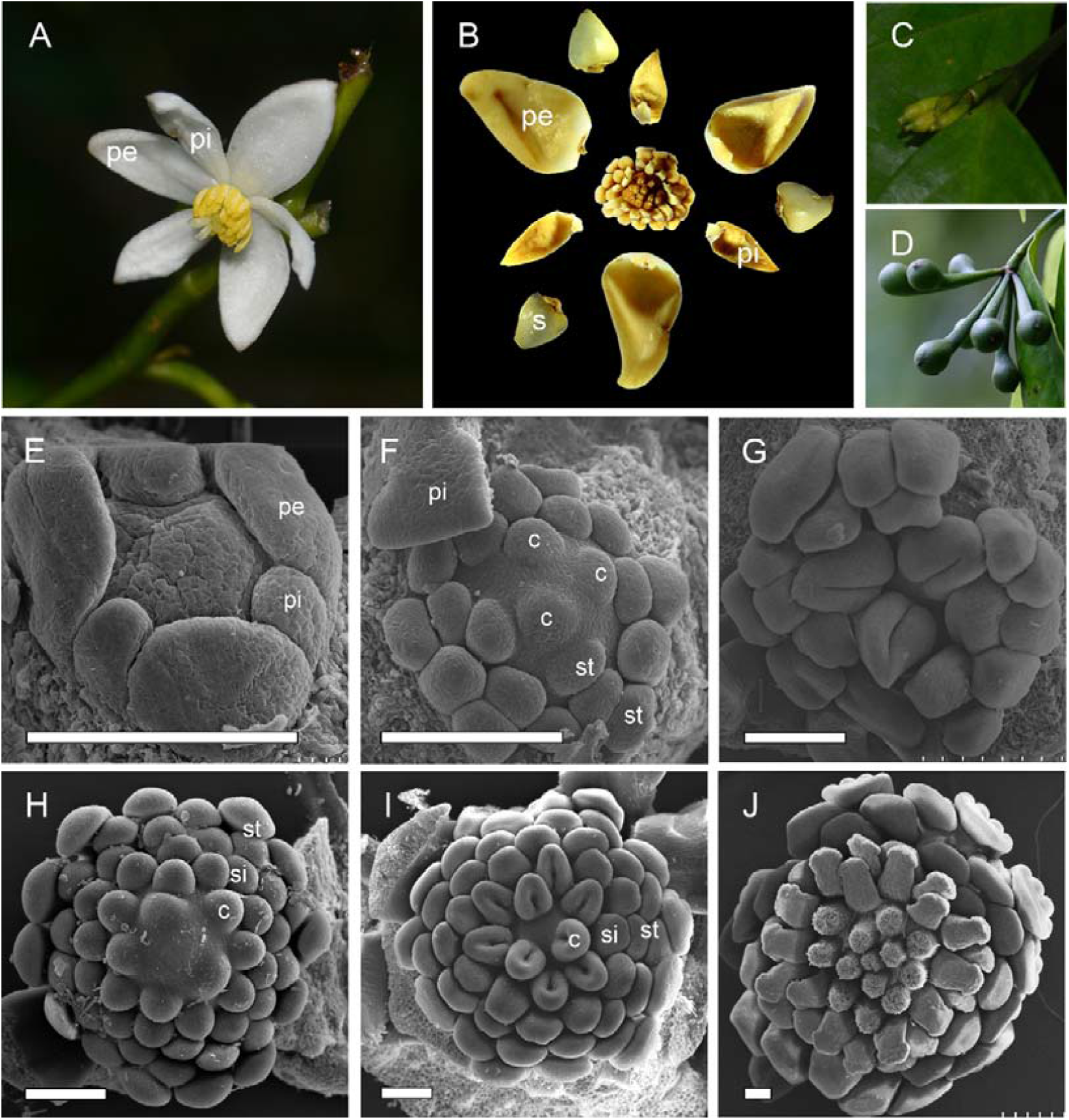
Floral morphology and gynoecium development in two *Anaxagorea* species. (**A**) *Anaxagorea luzonensis* flower. (**B**) *Anaxagorea javanica* flower. (**C**) Young *A. luzonensis* fruit. (**D**) Mature *A. javanica* fruit. (**E–G**) *A. luzonensis* floral development. (**H–J**) *A. javanica* gynoecium development. s, sepal; pe, outer petal; pi, inner petal; st, stamen; si, staminode; c, carpel. Scale bars =200 μm.

**Fig. 2.**
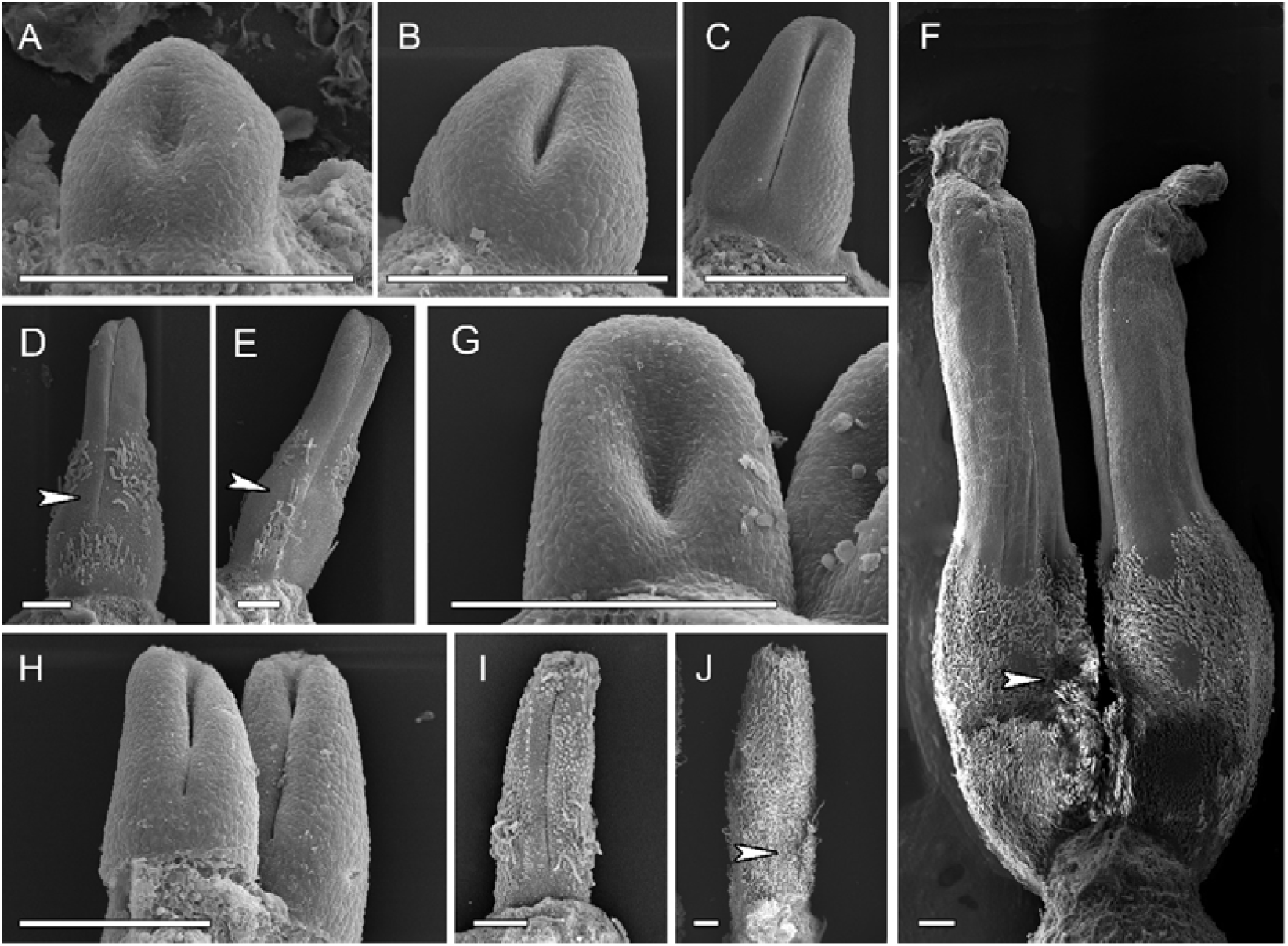
Carpel organogenesis in two *Anaxagorea* species. (**A–F**) *Anaxagorea luzonensis*. (**A**) Carpel primordia. (**B–C**) Carpel stipe emergence. (**D–E**) Carpel thickening and stigma formation, showing carpel stipe elongation. (**F**) Mature carpels. (**G–J**) *A. javanica* shows similar carpel developmental features to changes depicted in (**A–E**), and (**F**). Ventral slit end indicated by arrows. Scale bar = 200 μm.

**Fig. 3.**
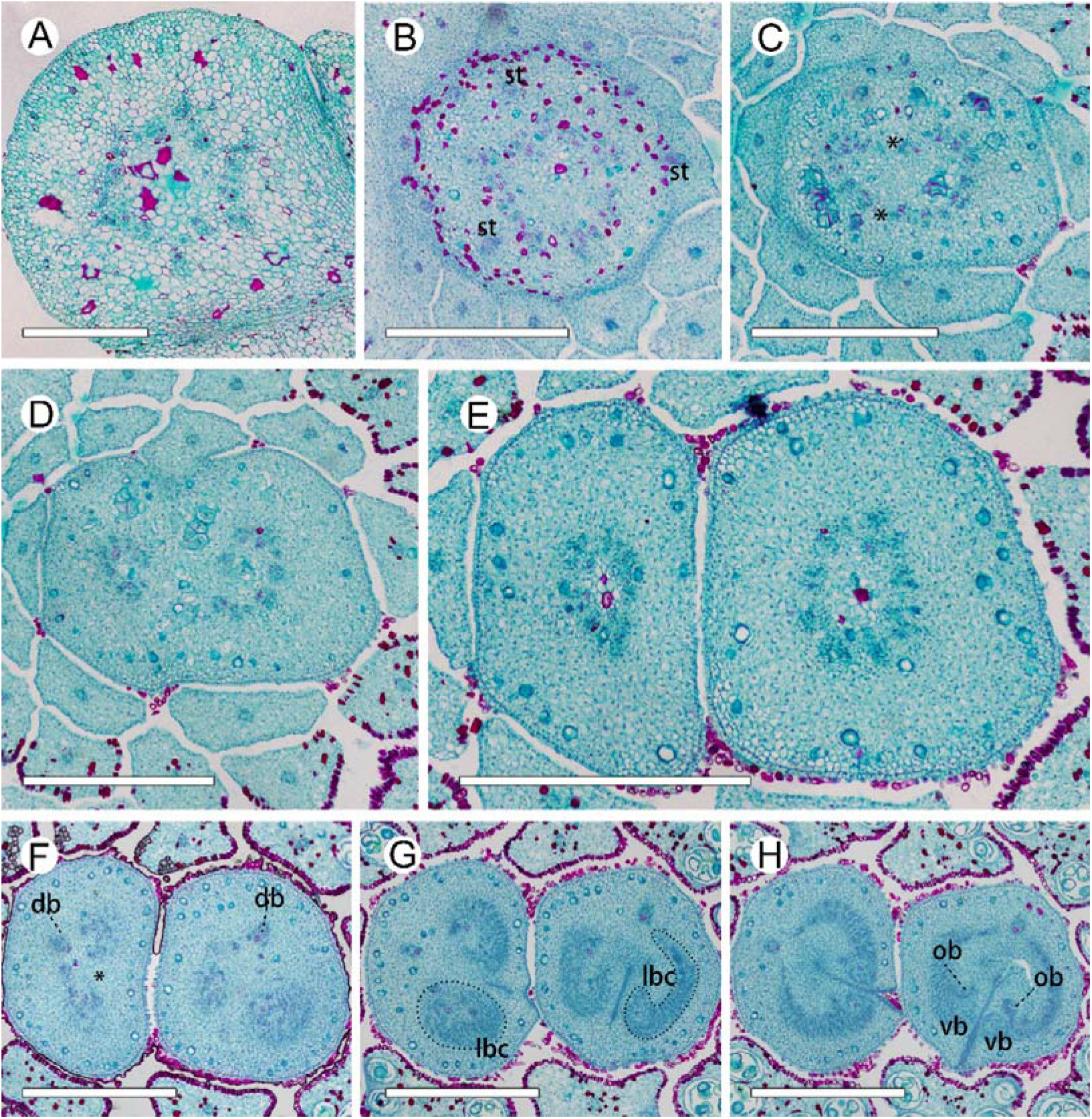
Ascending paraffin transections of the *Anaxagorea luzonensis* flower. (**A**) Base of receptacle. (**B**) Mid-section of androecia, showing stamen bundles and central stele. (**C**) Top of receptacle, showing central stele divided into two groups (* mark the breaks). (**D**) Bundles from the central stele enter carpels. (**E**) Base of carpels, showing basal ring. (**F**) Upper part of carpel stipes, showing the basal ring breaks (marked as *). (**G**) Bottom of ovary locule, showing amphicribral lateral bundle complexes (left), and “C”-shaped lateral bundle complexes (right). (**H**) Base of ovary locule. st, stamen; db, dorsal bundle; lbc, lateral bundle complex; vb, ventral bundle; ob, ovule bundle. Scale bar = 500 μm.

### Vasculature from receptacle to carpel

In the *A. luzonensis* cross-sections, the receptacle base presented a hexagon of 18 bundles from the pedicel stele (Fig. 3A). The hexagon had six breaks, which built up a crown of the cortical vascular system to supply the sepals and the two whorls of petals and the stamens (Figs. 3B). The central stele, composed of 18 bundles, finally broke into two nine-bundle groups at the floral apex and ran into the two-carpel gynoecium (Figs. 3C, 3D). Each group of nine bundles assembled as a basal ring around the parenchyma at each carpel base (Figs. 3E). At the slightly upper part of each carpel, several bundles emerged on the lateral side, and the basal ring broke, from which the dorsal bundle separated and the lateral bundles reorganized into two groups of lateral bundle complexes (Figs. 3F). In each of the lateral bundle complexes, the adjacent bundles tended to join, assembling into an amphicribral pattern (Fig. 3G). Below each placenta, each of the amphicribral lateral bundle complexes transformed into a set of “C”-shaped lateral bundle complexes, from which the ovule bundles separated, while the other bundles ran into the ovary wall. There were no horizontal connections between the dorsal and other bundles (Fig. 3H).

The pseudosteles at the base of the *A. Javanica* receptacle were triangular, with *∼* 45 bundles. The outer six cortical traces were cylindrical and served the sepals and petals (Figs. 4A, 4B). At a slightly higher level, the androecial bundles emerged and served the stamens by repeated branching, and the staminode bundles emerged as a crown around the central stele (Fig. 4C). Before entering the gynoecium, the central stele enlarged and broke up into *∼* 70 bundles to supply the nine carpels, and each carpel was served by 7–10 bundles (Figs. 4D–4E). The vascular bundle arrangement was similar to ascending sections in *A. luzonensis*, with the basal ring and amphicribral lateral bundle complexes presented in each carpel (Figs. 4F–4H).

**Fig. 4.**
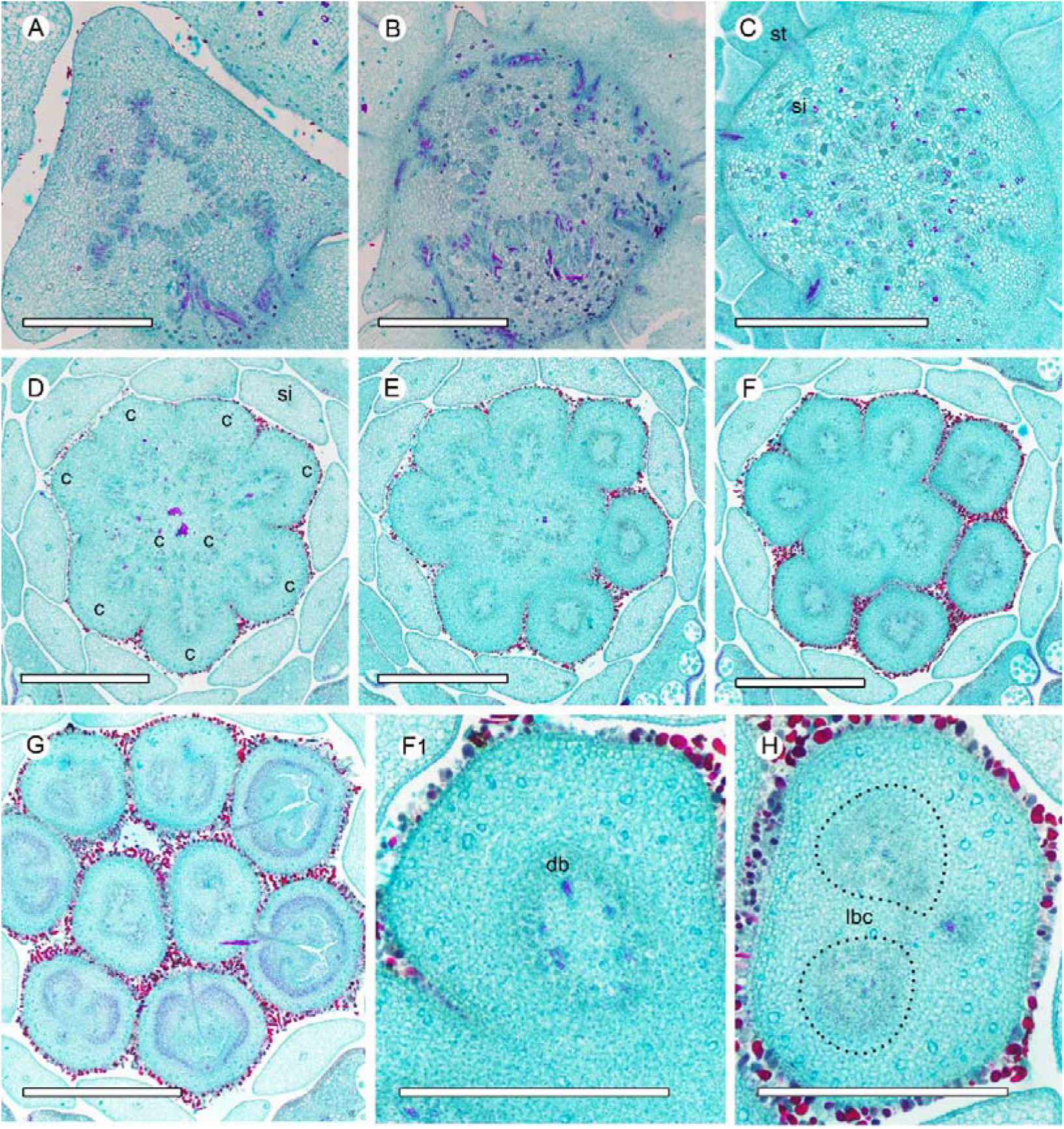
Ascending paraffin transections of the *Anaxagorea javanica* flower. **(A)** Base of receptacle, showing six groups of vascular bundles and sepal connections. **(B)** Points of petal connection to receptacle, showing perianth bundles. (**C**) Androecial bundles serving stamens by repeated branching. (**D–E**) Base of gynoecium, showing enlarged central stele breaks and bundles distributed into carpels. (**F–G**) Carpel vasculature at different positions. (**F1**) Detailed view of (**F**), showing basal ring of carpel. (**H**) Amphicribral lateral bundle complexes in carpel. st, stamen; si, staminode; c, carpel; db, dorsal bundle; lbc, lateral bundle complex. Scale bar = 500 μm.

### Three dimensional (3D)-reconstruction of carpel vascular topology

At the base of a mature *A. luzonensis* carpel, 15 discrete bundles were arranged in a radiosymmetrical pattern, forming a basal ring around the central parenchyma (Fig. 5A). At the slightly upper part, the basal ring curved inward on the ventral side and broke away from the invagination (Figs. 5B, 5C). The bundles (except the dorsal) divided into two groups on each side of the carpel, each forming a lateral bundle complex, which was also ring-arranged. At the flowering stage, the lateral bundle complexes corresponded to the above-mentioned sections of the amphicribral complexes (Figs. 5D–5F). Below each placenta, bundles of each lateral bundle complex broke up on the dorsal side and transformed into a “C”-shaped lateral bundle complex (Figs. 5G, 5H). The bundles on the ventral side of each lateral bundle complex gathered together (excluding the ventral bundle) and entered each ovule, while other bundles entered into the ovary wall. The ovule bundles were amphicribral. (Figs. 5I–5L).

**Fig. 5.**
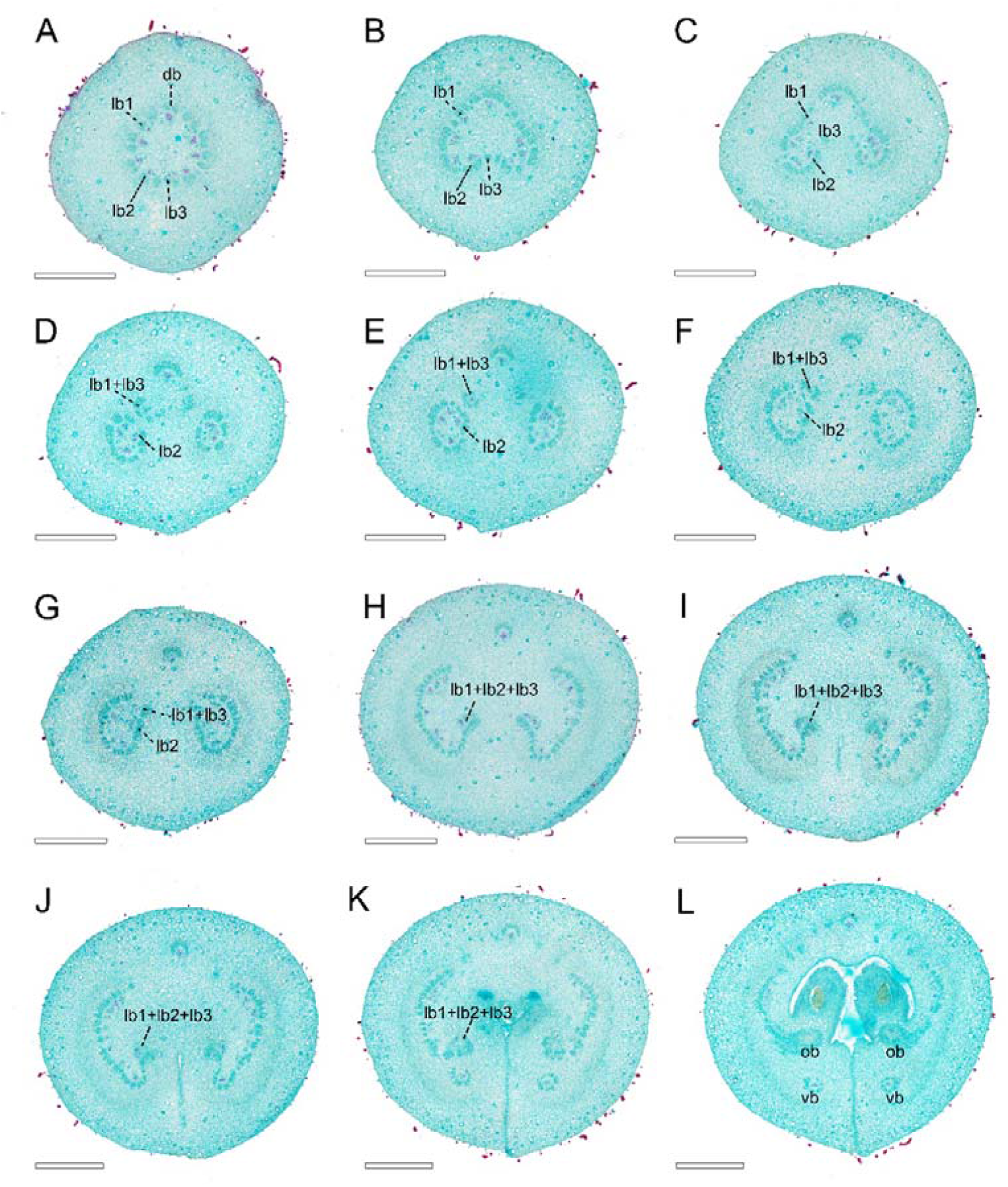
Ascending paraffin transections of the mature *Anaxagorea luzonensis* carpel. (**A**) Carpel base, showing basal ring. (**B–C**) Basal ring breaks on ventral side. (**D–F**) Ascending carpel stipe sections, showing lateral bundles reconstituted to two sets of ring-arranged lateral bundle complexes. (**G–H**) Top of carpel stipe, showing “C”-shaped lateral bundle complex. (**I–K**) Below ovary locule, showing formation of ovule bundles. (**L**) Base of ovary locule. db, dorsal bundle; lb, lateral bundle; vb, ventral bundle; ob, ovule bundle. Scale bar = 500 μm.

Consecutive cross-sections of *A. Javanica* were similar in vasculature to those of *A. luzonensis* (Figs. 6A–6D). The base of the mature *A. Javanica* carpel exhibited 16 distinct bundles forming the basal ring (Fig. 6A, 6F). The 3D model showed that (1) the basal ring and lateral bundle complex were cylindrical (Figs. 6F, 6H). (2) The ovules were fed directly by bundles from the base of the carpel through the lateral bundle complex. (3) Each ovule bundle was formed from several non-adjacent lateral bundles, and two bundles of them that fed each ovule joined on the ventral side (Figs. 6G, 6I). (4) The dorsal bundle remained independent throughout ontogenesis, without any link to other bundles (Appendix S1, S2; see Supplemental Data with this article).

**Fig. 6.**
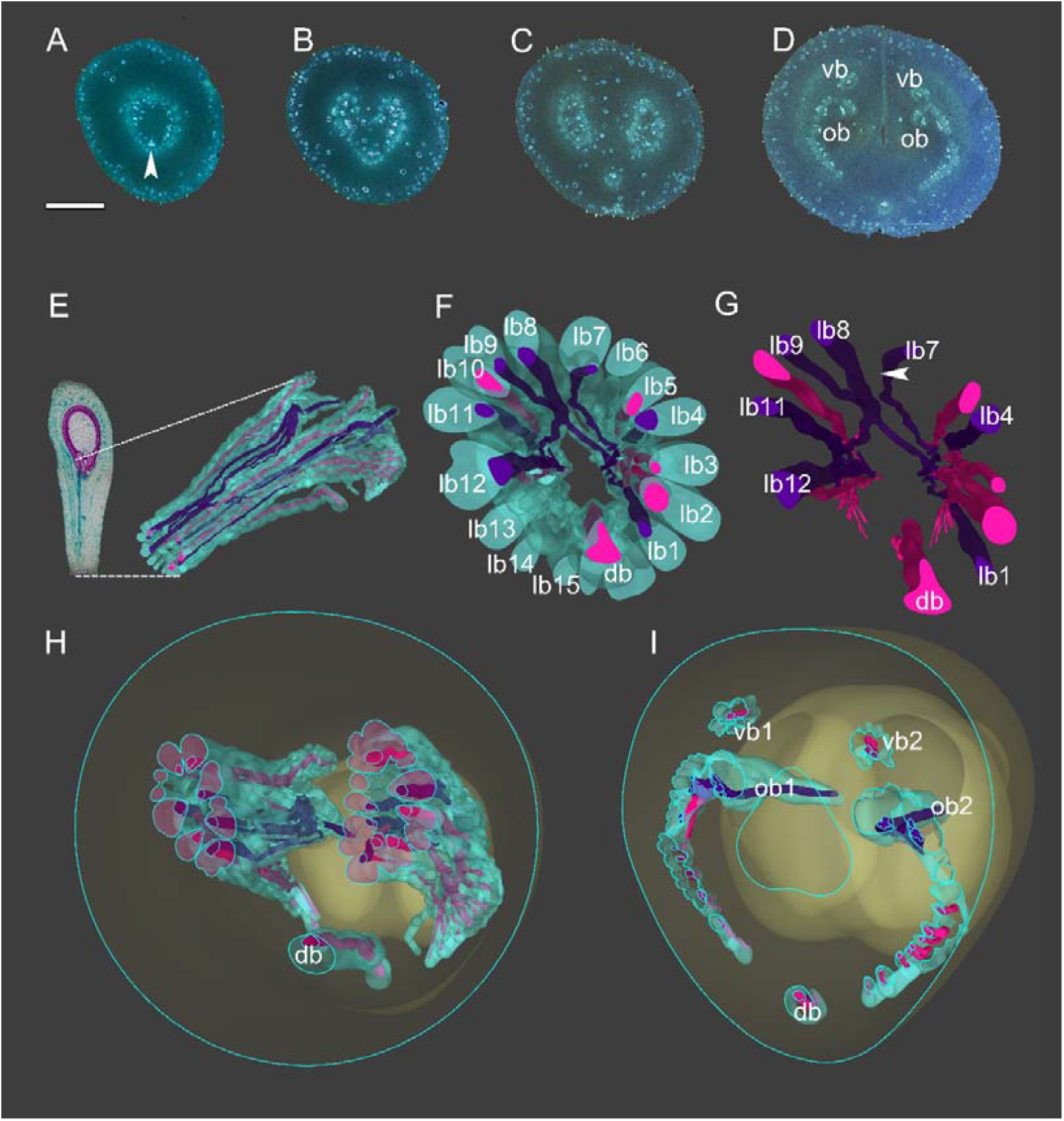
3D construction of *Anaxagorea javanica* vasculature. Bundle outlines colored green, xylem red, and purple, among which bundles associated with ovule bundles are colored purple. (**A–D**) Aniline blue-stained *A. javanica* sections for modeling. (**E**) Longitudinal section of mature *A. javanica* carpel (left), and 3D vasculature model, dotted lines on longitudinal section indicate vasculature position in carpel. (**F**) Perspective from base of carpel vasculature. (**G**) Perspective from base of carpel (xylem only). The arrow indicates the intersection of two lateral bundles which feed two ovules respectively. (**H**) Cross-section of 3D model corresponding to (**C**), showing ring-arranged lateral bundle complexes. (**I**) 3D model section showing distribution of vascular bundles at base of ovary. db, dorsal bundle; vb, ventral bundle; ob, ovule bundle, lb, lateral bundle. Scale bar = 500 μm.

## DISCUSSION

Elucidating the origin of carpel has been a challenge in botany for a long time (Takhtajan, 1969; Doyle, 1978; Cronquist, 1979; Thorne, 1996; Hickey & Taylor, 1996; Doyle, 2008). The carpel has been assumed as a conduplicate leaf-like structure bearing marginal ovules in the traditional interpretation. However, research was carried out to investigate whether the reproductive module (ovule/placenta) in the carpel and the carpel wall had evolved from two independent organs (Satina & Blakeslee 1941, 1943; Satina 1944, 1945; Angenent et al., 1995; Roe et al., 1997; Wynn et al., 2014; Zhang et al., 2019). Many hypotheses exist where the ovule/placenta of the carpel originated from the female reproductive branches of gymnosperms or seed ferns (e.g., the Gonophyll Theory [Melville, 1960]; the Neo-pseudanthial Theory [Doyle, 1994; Nixon et al., 1994]; the *Caytonia*-Glossopterid Model [Doyle, 2006]; the Unifying Theory [Wang 2010, 2018]). A logical inference from these hypotheses is that the placenta in angiosperms should have amphicribral vascular bundle, just like that in a young branch (Liu et al., 2014). Favoring this inference, amphicribral bundles have been documented in the placenta of many angiosperms (e.g., *Papaver* [Kapoor, 1973], *Psoraleae* [Lersten & Don, 1966], *Drimys* [Tucker, 1975], *Nicotiana* [Dave et al., 1981], *Whytockia* [Wang & Pan, 1998], *Pachysandra* [Von Balthazar & Endress, 2002], *Actinidia* [Guo et al., 2013], *Magnolia* [Liu et al., 2014]; *Michelia* [Zhang et al., 2017]; *Dianthus* [Guo et al., 2017]; *Tapiscia* [Xin et al., 2019]). However, regarding the amphicribral bundle as the evidence to support the axial nature of the carpel is considered to be flawed. As Endress (2019) summarized: (1) a vascular bundle often changes its anatomy along its length; the anatomy is influenced by its immediately adjacent histological environment; (2) the structure of a vascular bundle is not organ-specific; thus, it cannot be used to determine the kind of organ in which it was formed; and, (3) a vascular bundle is connected to other vascular bundles in the plant body, and its histology is greatly influenced by the next older vascular bundle to which it connects.

Considering the variability of the vascular bundles mentioned above, it is necessary to take the histological environment around the amphicribral ovule/placenta bundle into account. On the basis of confirming the existence of amphicribral ovule/placenta bundles in the carpel of *A. luzonensis* and *A. javanica*, this study also found that, vascular bundles are arranged in rings in different parts of the *Anaxagorea* carpel: (1) the basal ring maintains spatiotemporal continuity throughout the carpel stipe, and (2) the ring-arranged lateral bundle complexes at each placenta; these ring-arranged lateral bundle complexes developed from the amphicribral bundle complexes in the placenta with carpel maturation.

In most groups of Annonaceae, gynoecium is fed by an enlarged central stele, and each carpel is usually fed by three bundles, one median and two lateral (Deroin, 1989; Ronse De Craene, 1993; Deroin & Norman, 2016; Deroin & Bidault, 2017). This feature allows the carpel to be interpreted as foliar organ. Because in eudicots, usually one to three bundles connect the leaf with the vascular system of the stem (Beck, 2010). However, in *Anaxagorea*, the basal ring seems inconsistent with the traditional interpretation. In other sister branches, *Cananga* represents the relatively basal position. In the carpel stipe of *Cananga*, although three xylem bundles can be identified, the phloem of the vascular bundle is connected together (Appendix S3; see Supplemental Data with this article). This seems to represent a transitional pattern between *Anaxagorea* and other latter divided clades.

It has been reported that in *Anaxagorea*, the ovules are served by the lateral bundle complex from the base of the carpel [e.g., *A. luzonensis* (Deroin, 1997); *A. crassipetala* (Endress & Armstrong, 2011)]. In our study, the 3D model showed that the topological structure of the ring-arranged lateral bundle complexes plays a key role in forming the ovule bundles and that it facilitates the non-adjacent bundles to approach each other and merge. The dorsal bundle remained independent throughout, and there were no horizontal connections between dorsal bundle and the lateral bundle complexes. The ventral bundle participated in the formation of the spatial ring-arrangement of the lateral bundle complexes; however, it was not involved in forming of ovule bundles. In the basal ring, there are two lateral bundles which are fed to both ovules, which makes the topological structure of the basal ring unable to be flattened into a leaf-like structure bearing marginal ovules.

In conclusion, (1) the ovule/placenta bundles of *Anaxagorea* are amphicribral, (2) the ovule/placenta bundles are separated from the bundles of the carpel wall, and, (3) radiosymmetrical vasculature (including amphicribral bundles and ring-arranged bundle complexes) in the carpel of *Anaxagorea* are fed by a larger radiosymmetrical bundle system. Radiosymmetrical vasculature is a universal feature in vascular plant stems (Metcalfe & Chalk, 1979; Evert, 2006; Beck, 2010; McKown & Dengler, 2010; Evert & Eichhorn, 2013). Recent results show that a similar pattern of ovule/placenta bundles might be common in angiosperms. It is plausible that the carpel is the product of the fusion between an ovule-bearing axis and the phyllome that subtends it, and the placenta bundles originate from a former branch. However, comparable information on the ring-arranged bundles of carpels is still lacking. It is necessary for further investigation to understand whether the radiosymmetrical vasculature are universal in angiosperm carpel.

## ACKNOWLEDGMENTS

The authors thanks Profs. Lars Chatrou, Xin Wang, and Xin Zhang for their helpful suggestions to the manuscript. We also thank Chun-Hui Wang, Shi-Rui Gan, and Yan-Lian Qiu for their help in searching for the target species in the field, Qing-Long Wang for his assistance in caring for the transplant materials in Hainan province, Qiang Liu for his support during sampling, and Lan-Jie Huang and Ke Li for their advice on manuscript writing. We would also like to thank Editage for English language editing.

## FUNDING

This study was supported by the National Natural Science Foundation of China (grant number 31270282, 31970250).

## AUTHOR CONTRIBUTIONS

YL planned and designed the research, performed the experiments, collected the images, drew the illustrations, and wrote the article; WD performed the experiments and complemented the writing; YC developed the 3D model; SW complemented the writing; X-FW supervised the experiments and complemented the writing.

## SUPPORTING INFORMATION

Additional supporting information may be found online in the Supporting Information section at the end of the article.

S1 Serial Section-Based Carpel Vasculature 3D Reconstruction of *A. javanica* (3ds file)

S2 Serial Section-Based Carpel Vasculature 3D Reconstruction of *A. javanica* (movie)

S3 Ascending paraffin transections of *Cananga odorata* flower.

---

**Appendix S3.**
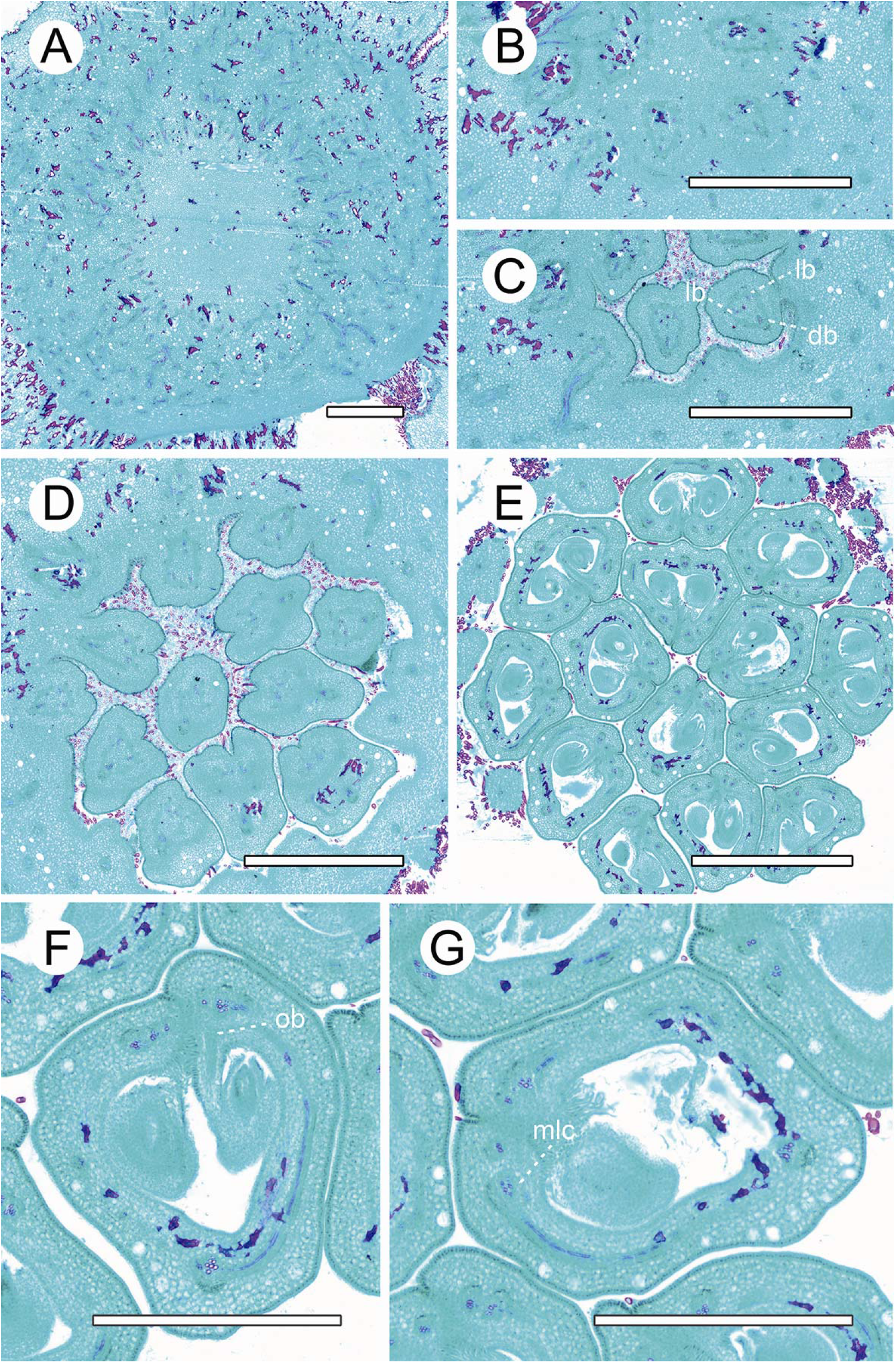
Ascending paraffin transections of the *Cananga odorata* flower. (**A**) Bundles of the receptacle. (**B**) The base of gynoecium. (**C**) The base of carpel, showing the connected phloem. (**D**) The sunken carpel stipes. (**E–F**) The ovary locule, showing ovules are fed by the dorsal bundle. db, dorsal bundle; lb, lateral bundle; mlc, carpel mediolateral bundle. Scale bars (**A–E**) = 1000 μm; **F, G** = 500 μm.

## Notes

### Competing Interest Statement

The authors have declared no competing interest.

## REFERENCES

Angenent, G. C., Franken, J., Busscher, M., van Dijken, A., van Went, J. L., Dons, H. J. & van Tunen, A. J. (1995). A novel class of MADS box genes is involved in ovule development in Petunia. Plant Cell 7, 1569–1582.

Beck, C. B. (2010). An introduction to plant structure and development: plant anatomy for the twenty-first century, 2nd edn. Cambridge, New York: Cambridge University Press.

Chatrou, L. W., Pirie, M. D., Erkens, R. H. J., Couvreur, T. L. P., Neubig, K.M., Abbott, J. R., Mols, J. B., Maas, J. W., Saunders, R. M. K. & Chase M.W. (2012). A new subfamilial and tribal classification of the pantropical flowering plant family Annonaceae informed by molecular phylogenetics. Botanical Journal of the Linnean Society 169, 5–40.

Chatrou, L. W., Turner, I. M., Klitgaard, B. B., Maas, P. J. M. & Utteridge, T. M. A. (2018). A linear sequence to facilitate curation of herbarium specimens of Annonaceae. Kew Bulletin 73, 39.

Cronquist, A. (1979). The evolution and classification of flowering plants. Brittonia 31(2), 293–293.

Dave, Y. S., Patel, N. D. & Rao, K. S. (1981). Structural design of the developing fruit of Nicotiana tabacum. Phyton 21, 63–71.

De Bary, A. & Scott, D. H. (1884). Comparative anatomy of the vegetative Organs of the phanerogams and ferns. Oxford, UK: Clarendon.

Ronse De Craene, L.P. & Smets, E. F. (1993). The distribution and systematic relevance of the androecial character polymery. Botanical journal of the Linnean Society 113, 285–350.

Deroin, T & Norman, É.M. (2016). Notes on the floral anatomy of Deeringothamnus Small (Annonaceae): cortical vascular systems in a chaotic pattern. Modern Phytomorphology 9, 3–12.

Deroin, T. (1988). Aspects anatomiques et biologiques de la fleur des Annonacées. Ph.D. dissertation, Centre d’Orsay, France: Université de Paris-Sud.

Deroin, T. (1989). Définition et signification phylogénique des systèmes corticaux floraux: l’exemple des Annonacées. In Comptes Rendus de l’Académie des Sciences Série III, Sciences de la vie, pp. 71–75. Paris, France: Elsevier.

Deroin, T. (1997). Confirmation and origin of the paracarpy in Annonaceae, with comments on some methodological aspects. Candollea 52, 45–58.

Deroin, T. & Bidault, E. (2017). Floral anatomy of Pseudartabotrys Pellegrin (Annonaceae), a monospecific genus endemic to Gabon. Adansonia 39, 111–123.

Doyle J. A. Origin of the angiosperm flower—A phylogenetic perspective. In Early evolution of flowers, Endress, P.K. & Friis, E.M. (eds.), Plant Systematics and Evolution Supplement 8, vol 8. Austria, Vienna: Springer.

Doyle J. A. (2006). Seed ferns and the origin of angiosperms. Journal of the Torrey Botanical Society 133,169–209.

Doyle, J. A. (1978). Origin of angiosperms. Annual Review of Ecology and Systematics 9, 365–392.

Doyle, J. A. (2008). Integrating molecular phylogenetic and paleobotanical evidence on origin of the flower. International Journal of Plant Sciences 169, 816–843.

Doyle, J. A. & Le Thomas, A. (1996). Phylogenetic analysis and character evolution in Annonaceae. Adansonia 18, 279–334.

Doyle, J. A., Sauquet, H., Scharaschkin, T. & Le Thomas, A. (2004). Phylogeny, molecular and fossil dating, and biogeographic history of Annonaceae and Myristicaceae (Magnoliales). International Journal of Plant Sciences 165, S55–S67.

Endress, P. K. (2019). The morphological relationship between carpels and ovules in angiosperms: pitfalls of morphological interpretation. Botanical Journal of the Linnean Society 189, 201–227.

Endress, P. K. & Armstrong, J. E. (2011). Floral development and floral phyllotaxis in Anaxagorea (Annonaceae). Annals of Botany 108, 835–845.

Esau, K. (1965). Plant anatomy, 2nd edn. New York, USA: Wiley.

Evert, R. F. (2006). Esau’s Plant Anatomy. Meristems, Cells, and Tissues of the Plant Body: Their Structure, Function, and Development, 3rd edition. New Jersy, USA: John Wiley & Sons, Inc, Hoboken.

Evert, R. F. & Eichhorn, S. E. (2013). Raven Biology of Plants, 8th edition. New York, USA: Palgrave Macmillan: W.H. Freeman Press.

Fahn, A. (1990). Plant Anatomy, 4th edition. Oxford, UK: Pergamon Press.

Guo, X. M., Xiao, X., Wang, G. X. & Gao R. F. (2013). Vascular anatomy of kiwi fruit and its implications for the origin of carpels. Frontiers in Plant Science 4, 391.

Guo, X. M., Yu, Y. Y., Bai, L. & Gao R. F. (2017). Dianthus chinensis L.: the structural difference between vascular bundles in the placenta and ovary wall suggests their different origin. Frontiers in Plant Science 8, 1986.

Hickey, L. J. & Taylor, D. W. (1996). Origin of the angiosperm flower. In Flowering Plant Origin, Evolution & Phylogeny, D.W. Taylor & L. J. Hickey (eds.), pp. 176–231. New York, USA: International Thomson publishing.

Kapoor, L. D. (1973). Constitution of amphicribral vascular bundles in capsule of Papaver somniferum. Linn. Botanical Gazette 134, 161–165.

Lersten, N. R. & Don, K. W. (1966). The discontinuity plate, a definitive floral characteristic of the Psoraleae (Leguminosae). American Journal of Botany 53, 548–555.

Liu, W. Z., Hilu, K. & Wang, Y. L. (2014). From leaf and branch into a flower: Magnolia tells the story. Botanical Studies 55, 28.

McKown, A. D. & Dengler, N. G. (2010). Vein patterning and evolution in C4 plants. Botany 88, 775–786.

Melville, R. (1960). A new theory of the angiosperm flower. Nature 188, 14–18.

Metcalfe, C. R. & Chalk, L. (1979). Anatomy of the dicotyledons, vol. 1, systematic anatomy of leaf and stem, with a brief history of the subject, 2nd edition. Oxford, UK: Clarendon Press.

Nixon, K. C., Crepet, W. L., Stevenson, D. & Friis, E. M. (1994). A reevaluation of seed plant phylogeny. Annals of the Missouri Botanical Garden 81, 484–533.

Ogura, Y. (1972). Comparative anatomy of vegetative organs of the pteridophytes. Berlin. Germany: Gebrueder Borntaeger.

Roe, J. L., Nemhauser, J. L. & Zambryski, P. C. (1997). TOUSLED participates in apical tissue formation during gynoecium development in Arabidopsis. Plant Cell 9, 335–353.

Rudall, P. J. (2007). Anatomy of flowering plants. An introduction to structure and development. Cambridge, UK: Cambridge University Press.

Satina, S. (1944). Perclinal chimeras in Datura in relation to development and structure (A) of the style and stigma (B) of calyx and corolla. American Journal of Botany 31, 493–502.

Satina, S. & Blakeslee, A. F. (1943). Perclinal chimeras in Datura in relation to the development of the carpel. American Journal of Botany 30, 395–450.

Satina, S. & Blakeslee, A. F. (1941). Periclinal chimeras in Datura stramonium in relation to development of leaf and flower. American Journal of Botany 28, 862–871.

Satina, S. (1945). Periclinal chimeras in Datura in relation to the development and structure of the ovule. American Journal of Botany 32, 72–81.

Takhtajan, A. (1969). Flowering plants, origin and dispersal. Edinburgh, UK: Oliver & Boyd Ltd.

Thorne, R. F. (1996). The Least Specialized Angiosperms. In Flowering Plant Origin, Evolution & Phylogeny, Taylor, D.W. & Hickey, L.J. (eds.), pp.286–313. New York, USA: Chapman & Hall.

Tucker, S. C. (1975). Carpellary vasculature and the ovular vascular supply in Drimys. American Journal of Botany 62, 191–197.

Von Balthazar, M. & Endress, P. K. (2002). Reproductive structures and systematics of Buxaceae. Botanical Journal of the Linnean Society 140, 193–228.

Wang, X. (2010). The Dawn Angiosperms. Heidelberg, Germany: Springer.

Wang, X. (2018). The Dawn Angiosperms, 2nd edition. Heidelberg, Germany, Springer.

Wang, Y. Z. & Pan, K. Y. (1998). Comparative floral anatomy of Whytockia (Gesneriaceae) endemic to China. In Floristic Characteristics and Diversity of East Asian Plants, Zhang, A.L. & Wu, S.G. (eds.), pp. 352–366. Beijing, China: China Higher Education Press.

Wynn, A. N., Seaman, A. A., Jones, A. L. & Franks, R. G. (2014). Novel functional roles for PERIANTHIA and SEUSS during floral organ identity specification, floral meristem termination, and gynoecial development. Frontiers in Plant Science 5, 130.

Xin, G., Ni, X. & Liu, W. (2019). Anatomy and Development of Gynoecium in Tapiscia sinensis Oliv. and its Implications for the Origin of Carpels. Bangladesh Journal of Botany 48(4), 933–941.

Zhang, X., Liu, W. & Wang, X. (2017). How the ovules get enclosed in magnoliaceous carpels. PLoS One, 12, e0174955.

Zhang, X., zhang, Z. X. and Zhao, Z. (2019). Floral ontogeny of Illicium Lanceolatum (Schisandraceae) and its implications on carpel homology. phytotaxa 416, 200–210. doi: 10.11646/phytotaxa.416.3.1

